# Chitin and cuticle proteins form the cuticular layer in spinning duct of silk-spinning arthropods

**DOI:** 10.1101/2021.11.04.466704

**Authors:** Xin Wang, Xiaoqian Xie, Kang Xie, Qingsong Liu, Yi Li, Xiaoyin Tan, Haonan Dong, Xinning Li, Zhaoming Dong, Qingyou Xia, Ping Zhao

## Abstract

Chitin is found in the exoskeleton and peritrophic matrix of arthropods, but recent studies have also identified chitin in the spinning duct of silk-spinning arthropods. Here, we report the presence and function of chitin and cuticle proteins ASSCP1 and ASSCP2 in the spinning duct of silkworms. We show that chitin and these proteins are co-located in the cuticular layer of the spinning duct. Ultrastructural analysis indicates that the cuticular layer has a multilayer structure by layered stacking of the chitin laminae. After knocking down ASSCP1 and ASSCP2, the fine structure of this layer was disrupted, which had negative impacts on the mechanical properties of silk. This work clarifies the function of chitin in the spinning duct of silk-spinning arthropods. Chitin and cuticle proteins are the main components of the hard and rigid cuticular layer, providing the shearing stress during silk fibrillogenesis and regulating the final mechanical properties of silk.

## INTRODUCTION

Arthropods, which are characterized by their exoskeleton, segmental body, and paired joint appendages, are the most successful and diverse animal species in our living world. Arthropods have significant capabilities for rapid adaptation and colonization in a variety of environments, such as the ground, ocean, and sky. The success of arthropods can be attributed to their unique characteristics. One of the features is the presence of the exoskeleton, which provides extensive protection from various harms, including mechanical damage, radiation, desiccation, and invasion by pathogenic microorganisms.

The exoskeleton is comprised of chitin and chitin-binding proteins (such as different types of cuticle proteins) (Liu, Zhang, & Zhu, 2019). Chitin is a linear polymer of the amino sugar N-acetyl-D-glucosamine, which can be easily found in the cuticle of the exoskeleton and the peritrophic matrix of the midgut in arthropods. It has been reported that the chitin content of the exuvial dry mass may be as high as 40% (Kramer, Hopkins, & Schaefer, 1995). The chitin content in the exoskeleton affects egg hatching (Zhu, Arakane, Beeman, Kramer, & Muthukrishnan, 2008), molting at various stages (He et al., 2013; Pesch, Riedel, Patil, Loch, & Behr, 2016), wing development (Chen, Yang, Tang, Yang, & Jin, 2017; Noh, Muthukrishnan, Kramer, & Arakane, 2018), and survival of arthropods (He et al., 2013; Su et al., 2016). In the peritrophic matrix, chitin is related to the permeability of small solutes and water (Agrawal et al., 2014), digestion of food (Liu et al., 2012), and size of the peritrophic matrix (Liu et al., 2012).

In addition to being present in the exoskeleton and peritrophic matrix, chitin is also found in the spinning duct of some silk-spinning arthropods, such as silkworms (Davies, Knight, & Vollrath, 2013), spiders (Davies et al., 2013), and caddisworms (Ashton & Stewart, 2019). The spinning duct of silk-spinning arthropods is the place where solid silk fiber is formed. It has been generally accepted that silk proteins undergo a gel-sol transition accompanied by physical and biochemical changes, such as pH (Askarieh et al., 2010; Hagn et al., 2010), metal ions (Wang, Li, Liu, Chen, et al., 2017; Wang, Li, et al., 2015; Wang, Zhao, et al., 2015; Zhou, Chen, Shao, Huang, & Knight, 2005), and shearing forces (Eisoldt, Hardy, Heim, & Scheibel, 2010; Holland, Urbach, & Blair, 2012; Rammensee, Slotta, Scheibel, & Bausch, 2008) in the spinning duct. Despite being an essential component of the spinning duct, the roles that chitin plays in this process remain unclear.

To understand the functions of the spinning duct in silk fibrillogenesis, RNA-seq and proteomics techniques have been introduced (Chang et al., 2015; Wang, Li, Liu, Xia, & Zhao, 2017; Wang et al., 2016; Yi et al., 2013). A large number of cuticle proteins have been identified in the silkworm spinning duct. Most of these proteins are predicted to have the chitin-binding domain (Wang, Li, Liu, Xia, et al., 2017; Yi et al., 2013). In our previous studies, we identified two cuticle proteins named ASSCP1 and ASSCP2 that are highly and specifically expressed in the spinning duct of silkworm (Wang et al., 2016; Yi et al., 2013). These proteins belong to the RR-2 family and possess the typical chitin-binding domain (Yi et al., 2013), which makes them the perfect targets to study the function of chitin in the spinning duct and the relationship between chitin and cuticle proteins.

In the present study, we aimed to investigate the functions of cuticle proteins and chitin in silk fibrillogenesis, as well as the expression pattern, localization, and chitin-binding activity of these proteins.

## MATERIALS AND METHODS

### Silkworms

Silkworms at different developmental stages were obtained from the Biological Science Research Center (Southwest University, China). Silk glands were dissected from the silkworm and divided into five parts (ASG, AMSG, MMSG, PMSG, and PSG) carefully according to their distinct morphology. After rinsing with sterilized water, all the samples were frozen rapidly in liquid nitrogen and stored at -80 ºC.

### Chitin content assay

Chitin content of the different parts of the silk gland was assayed according to the method of Zhang and Zhu (2006). Briefly, 0.1 g sample was homogenized with distilled water and centrifuged at 1800 ×*g* for 15 min at 25 ºC. The pellet was resuspended in 3% SDS. Next, the sample was incubated at 100 ºC for 15 min and centrifuged for 10 min. The resulting pellet was washed with distilled water, resuspended in 14 M KOH, and incubated at 130 ºC for 1 h to deacetylate chitin. The deacetylated sample was cooled on ice for 5 min and resuspended in ice-cold 75% ethanol. After being incubated on ice for 15 min, 35 μL of Celite 545 suspension was added to each sample. The samples were centrifuged at 1,800 ×*g* for 15 min at 4 ºC. The resulting pellet (insoluble chitosan) was washed with 40% cold ethanol and resuspended in distilled water. These samples were mixed with 10% NaNO_2_ and 10% KHSO_4_. After gentle shaking, samples were incubated at 25 ºC for 15 min to depolymerize the chitosan and deaminate the glucosamine residues from the chitosan. Samples were then centrifuged at 1,800 ×*g* for 15 min at 4 ºC and the supernatant was collected. After the addition of 12.5% NH_4_SO_3_NH_2_, the mixtures were vigorously shaken for 5 min at 25 ºC. Next, 0.5% freshly prepared 3-methyl-2-benzothiazolone hydrazine hydrochloride hydrate (MBTH) was added to each sample. The samples were incubated at 100 ºC for 5 min. After cooling for 10 min at 25 ºC, 20 μL of 0.83% FeCl_3_·6H_2_O solution was added, and the solution was thoroughly mixed. The cooled samples were placed on a 96-well microplate to determine the absorbance at 650 nm. Commercial glucosamine was used in a standard curve to determine the chitin content. All the chemicals and reagents were purchased from Sigma-Aldrich (USA).

### Western blotting

Proteins were extracted using 8 M urea with 30 mM DL-dithiothreitol (DTT). Protein concentration was determined using the method of Bradford. Samples (containing 20 μg of total protein) were mixed with loading buffer and separated on 12 % SDS-PAGE gels and transferred to PVDF membranes (Roche, Switzerland). After blocking with Tris-buffered saline with 0.1% Tween 20 and 5 % nonfat dry milk, the membrane was incubated with an affinity-purified specific antibody (1:10,000) and goat peroxidase-conjugated antirabbit IgG (1:20,000, Sigma-Aldrich, USA), for 2 h at 25 ºC. Immunoreactive protein signals were visualized using stable peroxide solution and luminol/enhancer solution (Millipore, USA). Silkworm α-tubulin protein was used as an internal control.

### Immunofluorescence

ASG samples were dissected on ice and fixed in 4% paraformaldehyde for 1 h at 25 ºC. Next, samples were frozen in Tissue-Tek (Sakura, Japan) at -80 ºC. Sections of 7 μm in thickness were cut by freezing microtome (Leica, Germany). The sections were used for immunofluorescence according to a previously described method (Wang, Zhao, et al., 2015). Chitin in the ASG was labeled using FITC conjugated wheat germ agglutinin (FITC-WGA, Sigma-Aldrich, USA).

### Expression and purification of recombinant protein

A fragment containing the RR-2 consensus sequence of *ASSCP1* (Fig. S1) was cloned and inserted into an expression vector (pET-28a). The primers used for PCR amplification are listed in Table S1. The vector was transformed into *Escherichia coli* BL21(DE3). The recombinant protein (ASSCP1-R) was induced by adding isopropyl β-D-1-thiogalactopyranoside (IPTG) into the LB media and purified by Ni-NTA affinity chromatography (Qiagen, USA). The full length of ASSCP2 protein (ASSCP2-R) was also expressed and purified according to the same protocols. Then, the rabbit polyclonal antibodies against ASSCP1 or ASSCP2 were produced and purified commercially by Zoonbio Biotech (China).

### Preparation of chitin-binding proteins in ASG

ASG samples were homogenized in 20 mM PBS buffer (pH 7.4) with 15 mM NaCl and centrifuged at 12,000 × *g* for 20 min at 4 ºC. Proteins were then extracted from the pellets in 20 mM PBS buffer (pH 7.4) containing 2 % SDS. The mixture was vortexed at 25 ºC for 2 h, centrifuged at 12,000 ×*g* for 20 min at 25 ºC, and the supernatant was collected. Next, 1 M KCl was added into each sample to remove most of the SDS. The supernatant was collected by centrifugation (12,000 ×*g*, 10 min, 4 ºC) and diluted 20-fold with 1 mM HEPES (pH 7.4). To remove residual SDS, an ultrafiltration tube (MWCO 10 kDa, Millipore) was used by adding 1 mM HEPES (pH 7.4) into the solution and centrifuging at 4,000 ×*g* for 20 min at 4 ºC repeatedly until the SDS concentration was sufficiently low (<0.001%). The concentration of SDS in the samples was determined according to a previously described method (Arand, Friedberg, & Oesch, 1992).

### Chitin and cellulose binding assay

Chitin and cellulose binding assay was performed as previously described (Tang, Liang, Zhan, Xiang, & He, 2010). Briefly, chitin (New England BioLabs, UK) and cellulose (Sigma-Aldrich, USA) beads were equilibrated in 20 mM HEPES (pH 7.4) containing 15 mM NaCl. A total of 3 mL solution (containing 3 mg ASG proteins or recombinant proteins) was mixed with 1 mL equilibrated chitin or cellulose beads for 3 h at 4 ºC with gentle shaking. Next, the slurries were packed into a column and washed with 20 mM HEPES (pH 7.4) and 1 M NaCl five times. The flow-through component and the washing eluate are referred to as F and W fractions, respectively. The bound proteins were eluted with 20 mM HEPES (pH 7.4) containing 15 mM NaCl and 8 M urea. The resulting eluate is referred to as E fraction. All these fractions were collected and used for subsequent SDS-PAGE and western blotting analyses.

### Transgenic vector construction and transgenic silkworm isolation

To study the physiological functions of ASSCP1 and ASSCP2, we constructed two transgenic vectors in which the dsRNA of *ASSCP1* or *ASSCP2* gene was controlled by a spinning duct specific promoter (*BmCP231*). The fragments of the dsRNA of the *ASSCP1* or *ASSCP2* gene were synthesized by Sangon Biotech Company (China) according to the CDS sequences of these genes. The sequences can be found in Table S2. In these vectors, the *RFP* gene or *GFP* gene acted as the marker gene that was driven by the *3×P3* promoter and expressed in the compound eyes and nervous tissues of *B. mori* (Fig. S5). All the primers used are listed in Table S1. After purification, each vector was microinjected into silkworm eggs. The generation of non-diapause embryos, microinjection, and screening of transgenic silkworms were performed as previously described (Wang, Zhao, et al., 2015). The transgenic silkworm line was named *ASSCP1-* and *ASSCP2*^*-*^, respectively. By cross-breeding, we obtained the transgenic line (ASSCP1^-^/ASSCP2^-^) that both ASSCP1 and ASSCP2 were knocked down. The economic trait (cocoon shell ratio) of transgenic silkworm lines was assessed by weighing the cocoon and the pupae. Total proteins from ASGs at the 4^th^ molting, 5^th^ instar, and spinning stages were extracted, and western blotting was used to determine whether the expressions of these two cuticle proteins were successfully knocked down in these developmental stages. Chitin content assay was also performed.

### Morphological analysis

Morphological analysis of ASG was performed using an EVOS FL Auto Microscope system (Life Technology, USA). All sections of ASG were observed under a confocal microscope (Olympus, Japan). The surface of the cocoon and all the fibers were observed using scanning electron microscopy (SEM, Hitachi, Japan) at an acceleration voltage of 5 kV and a polarizing optical microscope (Leica, Germany). For transmission electron microscope (TEM) observations, fresh ASG was dissected, cut into 1 mm^3^ section, and fixed in ice-cold 2.5 % glutaraldehyde solution (Sigma-Aldrich, USA) for 24 h. ASG sections were then dehydrated in an ethanol series from 50% to 90%, embedded in Spurr resin (Spi-Chem, USA), cured, and cut into 70-nm sections. Sections were stained using 3% uranyl acetate-lead citrate (Spi-Chem, USA). A TEM (JEM1230, JEOL, Japan) was used to capture the images. For the immunoelectron microscopy, resin blocks were cut to 70 nm thin and the tissues were fished out onto the 150 meshes nickel grids with formvar film. After rinsing, the tissues were blocked for 30 min in a TBS solution containing 1% BSA. Then, the nickel grids were incubated in the ASSCP1 antibody (1:30) diluted in TBS solution containing 1% BSA. After rinsing on TBS solution 3 times, the nickel grids were incubated in the secondary antibody solution and washed by TBS solution. 2% uranium acetate saturated alcohol solution was used for staining the nickel grids. After that, the nickel grids are observed under the TEM. The 10 nm black golden particles are positive signals of the target proteins.

### FTIR analysis of silk fibers

The degumming process, FTIR analysis, and peak deconvolution were performed according to the previous literatures (Peng et al., 2019; Wang, Li, Liu, Chen, et al., 2017). Briefly, five cocoons from each silkworm line were chosen randomly and degummed by boiling in two 30 min rounds using 0.5% (w/v) NaHCO_3_ solution. The degummed silk fibers were washed with distilled water and allowed to air dry at room temperature. Then, infrared spectroscopy in attenuated total internal reflection mode was performed on these fibers using a Thermo Scientific (USA) Nicolet iN10 with a Slide-On ATR objective lens. The spectra were recorded in the 650−4000 cm^−1^ range at a resolution of 8 cm^−1^ with 256 scans for each measurement. The applied ATR current pressure was set to 75. OMNIC 9 software (ThermoScientific, USA) was used to collect and process the spectral data. Baseline correction, deconvolution of amide I bands, and peak fitting were performed using OMNIC 9 software (ThermoScientific) and PeakFit software (Seasolve, version 4.12, USA). The content of each secondary structural component is determined by measuring the ratios of areas under the corresponding peaks (Fig. S10).

### Mechanical testing of silk fibers

To accurately compare the mechanical properties of silks between each silkworm line, we used the forcible silking method to obtain single silk fibers. Since the environmental conditions have a great influence on the mechanical properties of silks, it is easier to precisely control these conditions by forcible silking. Briefly, spinning silkworms were divided into weight-matched groups. The single silk was grasped and reeled artificially from the silkworm spinneret using a device developed by Khan et al. (Khan et al., 2008) and modified by ourselves. We attempted to obtain silk fibers at 60 rpm/min for more than 10 min. The single silk fiber was cut to a length of 10 mm. The diameter of each silk fiber was measured across two brins under optical microscopy (Zeiss, Germany). The fiber cross-section area was calculated by using the diameter. It should be noted that the cross-sectional shape of silk is approximately elliptical rather than circular, so the cross-sectional area of silk was overestimated in the calculation. Tensile tests were performed on a dynamic mechanical analyzer Q800 (DMA, TA Universal Analysis, USA) at 25 °C and 60% humidity. The stretching speed was set to 1 mm/min. Raw data were collected and analyzed using the TA Universal Analysis software. Subsequently, the data were used to calculate mechanical performance parameters using ORIGIN 8.0 (OriginLab, USA). Since the cross-sectional area of silk was overestimated, the stress of the silk fiber was underestimated accordingly.

## RESULTS

### ASSCP1 and ASSCP2 are co-located with chitin in the cuticular layer of the silkworm spinning duct

RNA-Seq and proteomics approaches have identified ASSCP1 and ASSCP2 as the silkworm spinning duct specific cuticle proteins, which are predicted to have the chitin-binding domain (Chang et al., 2015; Wang et al., 2016; Yi et al., 2013). If these proteins can bind chitin, they should be co-located with chitin in the same tissue. To test this possibility, western blotting and immunofluorescence were performed (Fig. 1). Anterior silk gland (ASG), spinneret, and Filippi’s gland constitute the spinning duct of the silkworm. Because the spinneret and Filippi’s gland are both quite small, we used the ASG to perform the experiments. Tissue distributions of ASSCP1 and ASSCP2 revealed that they were only expressed in ASG (Fig. 1A). Furthermore, the localization of these two proteins and chitin were determined using frozen sectioning and immunofluorescence. As shown in Fig. 1B, the green signal was detected in the cuticular layer of ASG, indicating that the chitin is located in this region. The signals from ASSCP1 and ASSCP2 were also detected in the cuticular layer (Fig. 1B), suggesting that both cuticle proteins and chitin are co-located in the cuticular layer of the silkworm spinning duct.

**Fig. 1.**
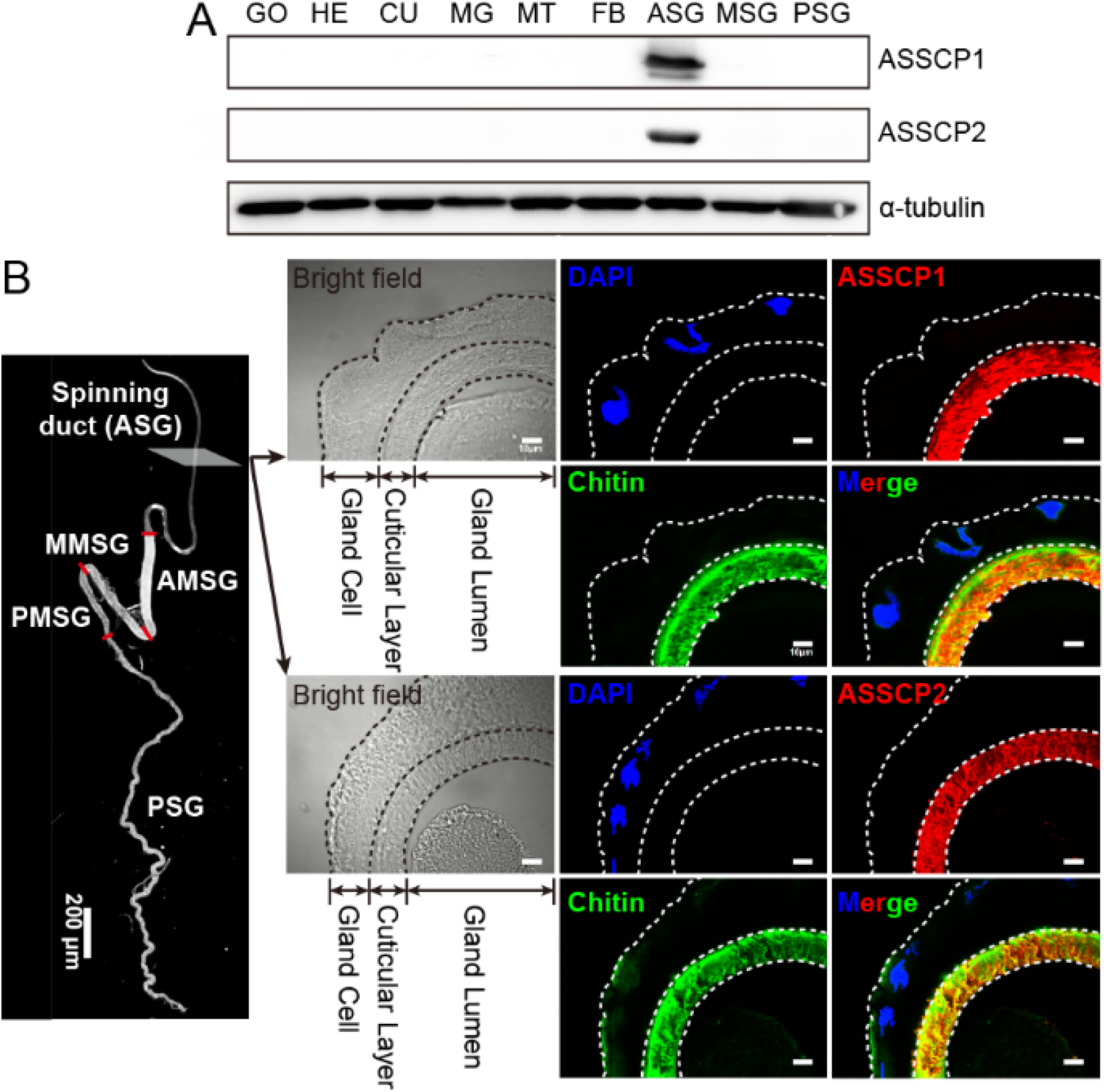
ASSCP1, ASSCP2, and chitin are co-located in the cuticular layer of the silkworm spinning duct. **(A)** Western blotting analysis of ASSCP1 and ASSCP2 in several tissues from the third day of fifth instar silkworm larvae. GO, gonad; HE, head; CU, cuticle; MG, mid-gut; MT, malpighian tubule; FB, fat body; ASG, anterior silk gland; MSG, middle silk gland; PSG, posterior silk gland. **(B)** Localizations of ASSCP1, ASSCP2, and chitin in the ASG from the spinning silkworm. Blue, the signal from the nucleus. Red, signals from ASSCP1 and ASSCP2. Green, the signal from chitin.

### ASSCP1 and ASSCP2 are chitin-binding proteins

Although chitin has been found in the spinning duct of silkworms (Davies et al., 2013), the exact content of chitin in this tissue is still unclear. In the present study, the distributions of chitin in different parts of the silk gland were analyzed. Compared with other parts of the silk gland, the chitin content in ASG is very high (Fig. 2A), whereas the chitin content in MSG and PSG is quite low. We have found that ASSCP1 and ASSCP2 are co-located with chitin in the cuticular layer of the spinning duct. To investigate whether these proteins could bind chitin, chitin-binding assays were performed using proteins extracted from ASG (AP, Fig. 2B). As shown in Fig. 2B, the protein bands were detected in the elution fractions (E’ and E’’). Using the ASSCP1 and ASSCP2 specific antibodies, we identified the band in E’ and E’’ fraction as ASSCP1 and ASSCP2, respectively. These results were further confirmed by matrix assisted laser desorption ionization-time of flight mass spectrometry (MALDI-TOF MS). Then, we successfully generated and purified a truncated ASSCP1 recombinant protein (ASSCP1-R) and a full-length ASSCP2 recombinant protein (ASSCP2-R) using an *E. coli* expression system (Figs. S1 and S2). The reason why we chose to express the truncated form of ASSCP1 protein is the repeat sequences (eg, SESSSEE, Fig. S1) in the C-terminus prevent expression of the full-length protein.

**Fig. 2.**
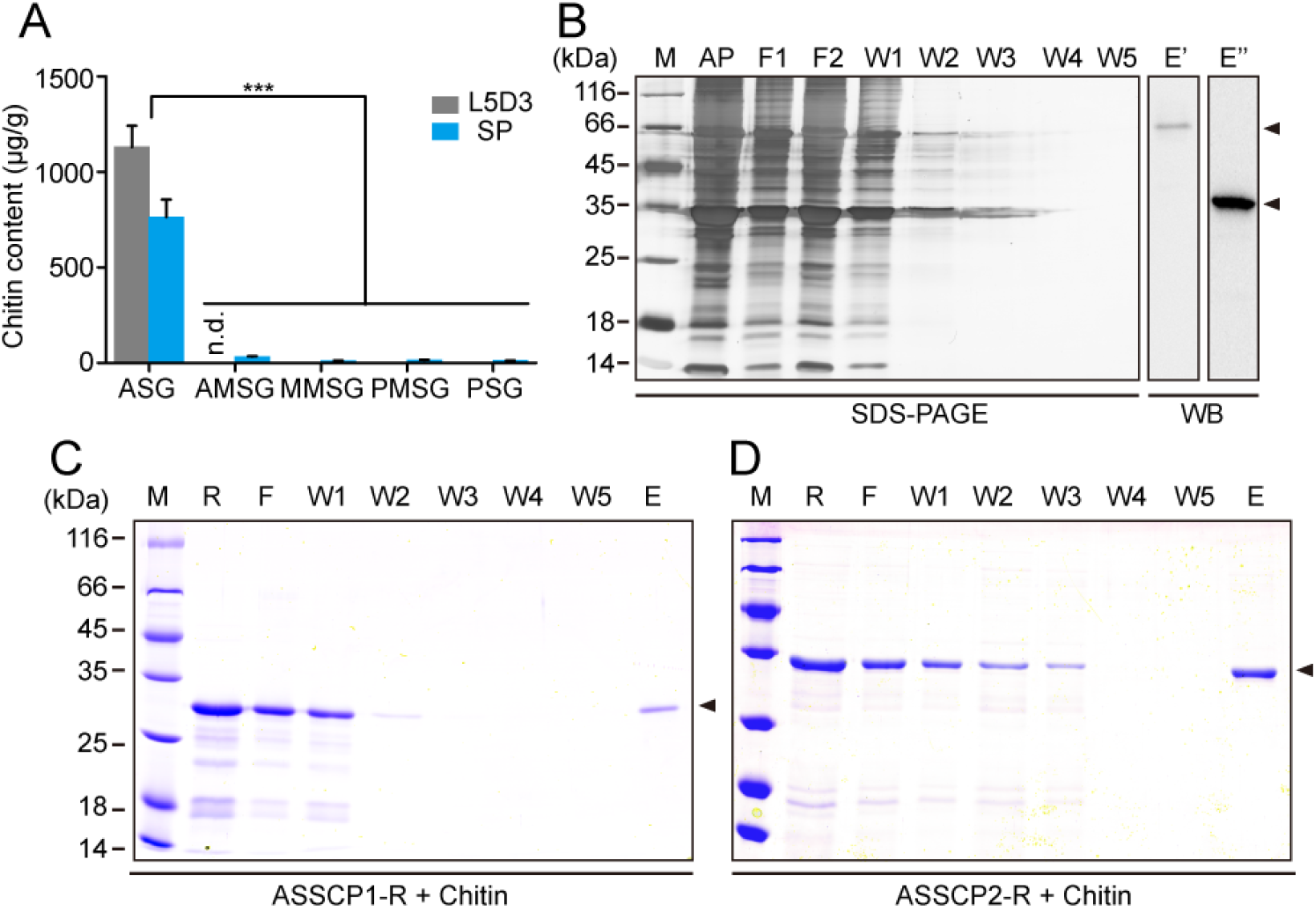
ASSCP1 and ASSCP2 are chitin-binding proteins. **(A)** Distributions of chitin in different parts of the silkworm silk gland. The decrease in chitin content at the spinning stage was due to the increase in the total weight of ASG. n. d., not detected; L5D3, the sample from the 3rd day of 5th instar larvae; SP, the sample from the spinning larvae; ASG, anterior silk gland; AMSG, the anterior section of middle silk gland; MMSG, the middle section of middle silk gland; PMSG, the posterior section of middle silk gland; PSG, posterior silk gland. ***, *p*-value < 0.001. **(B)** Chitin binding assay of ASG-extracted proteins. ASG proteins were extracted and incubated with commercial chitin beads. Non-specific bound proteins were washed with NaCl and the chitin-binding proteins were eluted with urea. Protein fractions were detected using SDS-PAGE followed by silver nitrate staining and western blotting (WB) with ASSCP1 and ASSCP2 specific antibodies, respectively. Gel condition, 12 % SDS-PAGE; M, protein marker; AP, ASG-extracted proteins; F1-F2, flow-through fractions; W1-W5, washing fractions; E’ and E’’, eluted fractions; Arrowhead in E’, ASSCP1 protein; Arrowhead in E’’, ASSCP2 protein. **(C)** Chitin binding assays of purified recombinant ASSCP1 protein (ASSCP1-R). All fractions were detected using SDS-PAGE followed by Coomassie brilliant blue G-250 staining. M, protein marker; R, recombinant protein; F, flow-through fraction; W1-W5, washing fractions; E, eluted fraction; Arrowhead, ASSCP1-R. **(D)** Chitin binding assays of purified recombinant ASSCP2 protein (ASSCP2-R). Arrowhead, ASSCP2-R.

Chitin and cellulose binding assays were performed using the recombinant proteins. Cellulose was used as a negative control. As shown in Fig. 2C and 2D, the signal was only detected in the elution fraction (E) after recombinant proteins were incubated with chitin beads. In contrast, no signal corresponding to cellulose binding was observed (Fig. S3). These results suggest that ASSCP1 and ASSCP2 are chitin-binding proteins

### Cuticle proteins and chitin are periodically present in the silkworm larval stage

Western blotting was performed to detect the presence of ASSCP1 in different instar larvae. Temporal expression analysis revealed that this protein is expressed only during larval feeding stages (Fig. 3A). The repetitive expression pattern of ASSCP1 during the silkworm feeding-molting transition led us to determine the exact time points of its synthesis and degradation. Thus, we further performed qRT-PCR and SDS-PAGE analysis using the ASG samples from different time points during the fourth molting larvae (Fig. 3B and 3C). It can be seen that the proteins had undergone degradation when larvae began to molt (0 h - 3 h). From 6 h to 12 h after molting, no positive protein bands could be detected but the transcription of the gene was active. The positive signals appeared again in the late molting stage (15 h after molting), indicating that the proteins are re-synthesized.

**Fig. 3.**
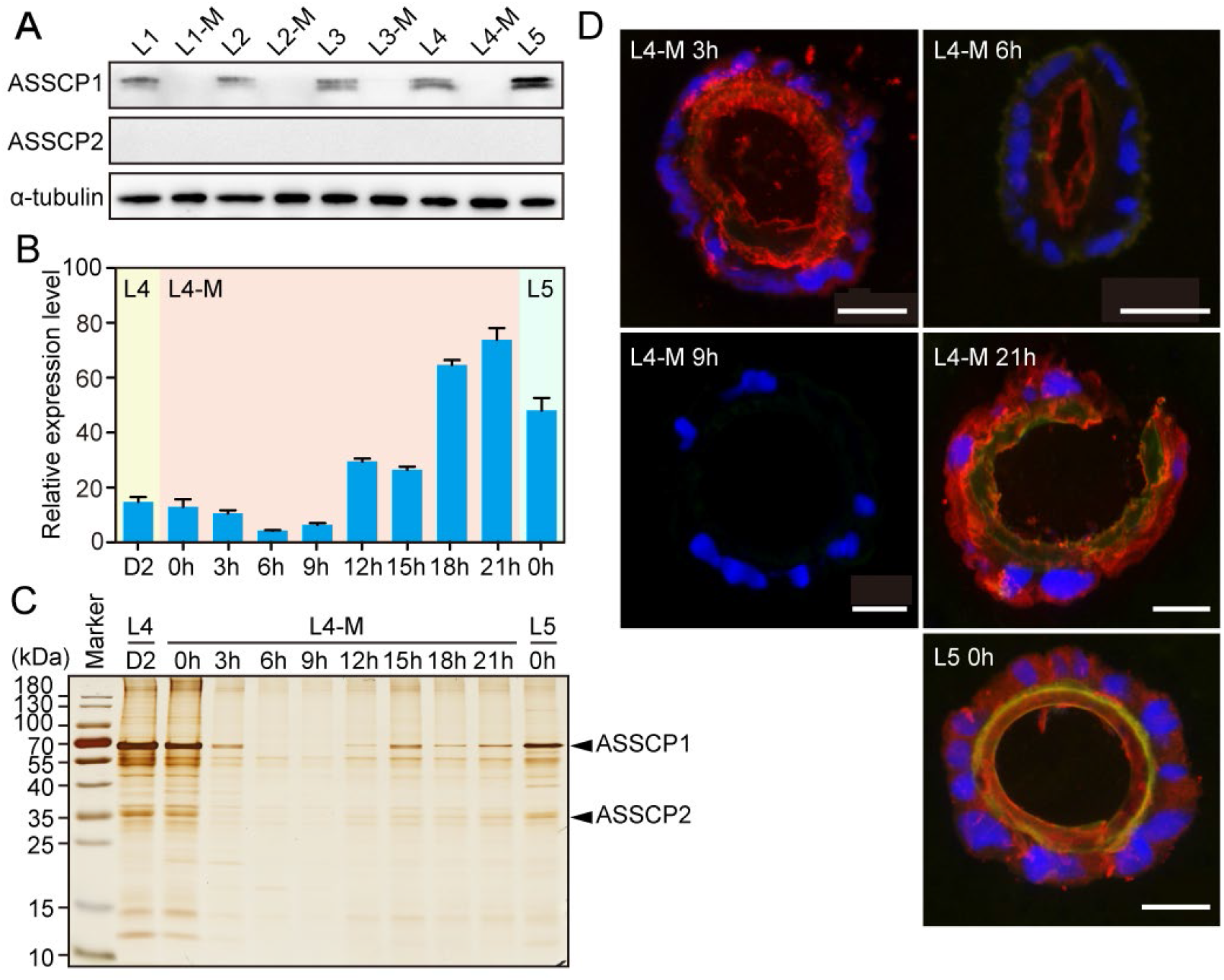
Cuticle proteins and chitin are periodically expressed. **(A)** Expression profile of ASSCP1 protein in the whole body of silkworm larvae from the first to fifth instar. The expression of ASSCP2 was quite low in these stages, no positive signal could be detected. L1, newly hatched larvae; L1-M, molting larvae on the first instar; L2, feeding larvae on the second instar; L2-M, molting larvae on the second instar; L3, feeding larvae on the third instar; L3-M, molting larvae on the third instar; L4, feeding larvae on the fourth instar; L4-M, molting larvae on the fourth instar; L5, feeding larvae on the fifth instar. **(B)** Expression profile of *ASSCP1* transcripts during the fourth molting stage using qRT-PCR. **(C)** Expression profile of ASSCP1 and ASSCP2 proteins during the fourth molting stage using SDS-PAGE, followed by silver nitrate staining. **(d)** Immunofluorescence localization of ASSCP1 in ASG during molting stage in the fourth instar. Blue, nucleus; Green, chitin; Red, ASSCP1 protein; Scale bar, 50 μm.

Immunofluorescence results clearly show that the cuticular layer was renewed during the IV molting stage (Fig. 3D). Consistent with western blotting results, the positive signal of ASSCP1 (red) and weak signal for chitin (green) were detected in the cuticular layer 3 h after molting. However, the shape of the cuticular layer was irregular, suggesting the degradation process had begun. Six hours after molting, only ASSCP1 could be detected and the cuticular layer was very thin. The cuticular layer was completely degraded 9 h after molting. At the late molting stage, ASSCP1 and chitin signals were both detected in the gland cell and cuticular layer, which indicates that the layer is re-built.

### Cuticle proteins and chitin are constantly present during silk spinning and degraded during pupation

Signals from cuticle proteins and chitin were constantly observed in the cuticular layer during the whole feeding stage of the fifth instar (Fig. S4). Assumed as an essential component of the spinning duct, chitin should be present during the silk spinning process. Thus, we measured the thickness of the cuticular layer during silk spinning to confirm this assumption. Fig. 4A shows that the thickness of the cuticular layer remained unchanged during the silk spinning process. However, the cuticular layer underwent a dramatic degradation during the pupation stage, when silk spinning was complete. We performed morphological analysis of the ASG and found that the ASG morphology remained unchanged during the silk spinning process, but the gland cell contracted and showed obvious apoptotic characteristics during pupation (Fig. 4B). Immunofluorescence analysis revealed that chitin and cuticle proteins were also constantly presented during silk spinning (Fig. 4C).

**Fig. 4.**
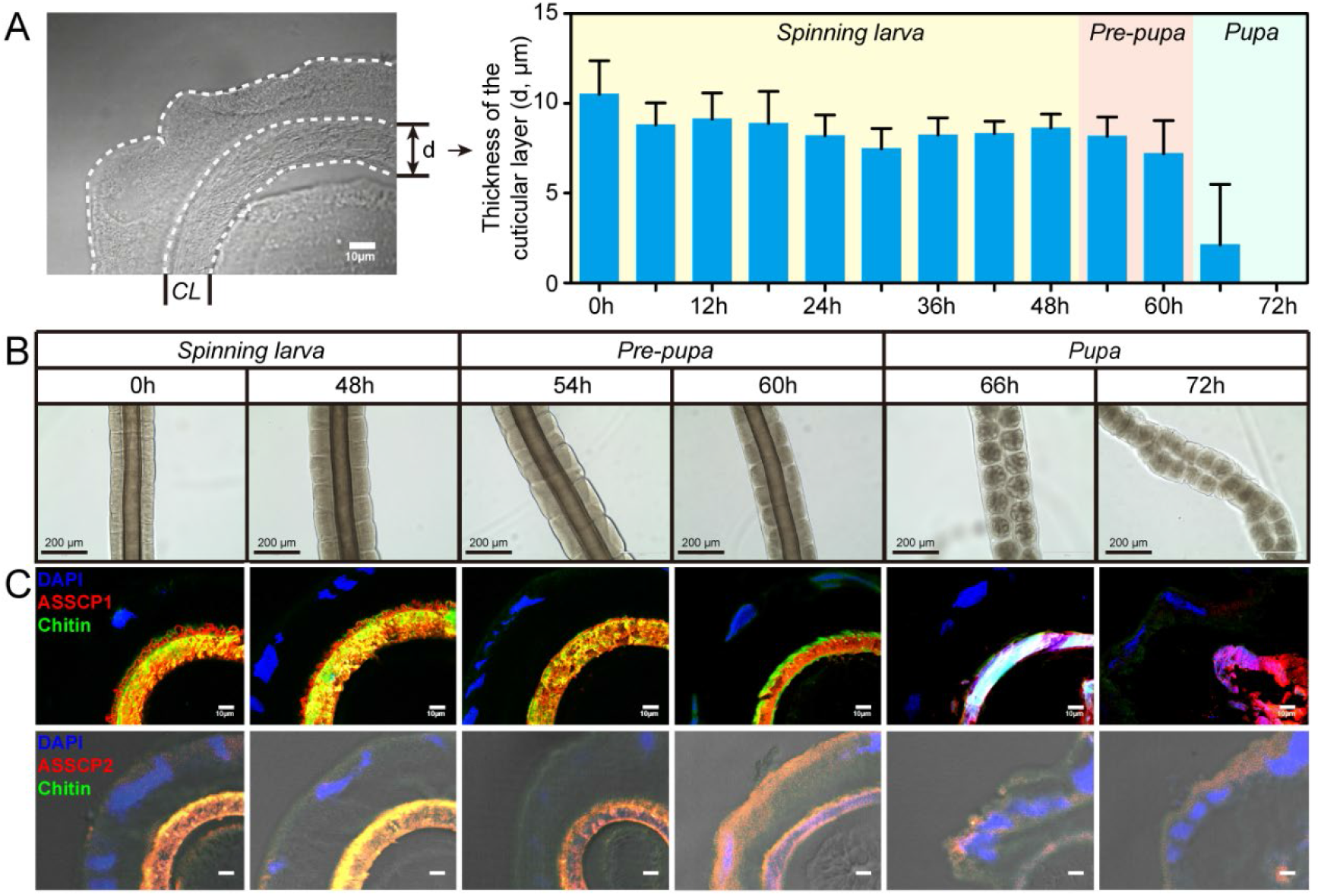
Cuticle proteins and chitin are constantly present during silk spinning and are degraded during pupation. **(A)** The thickness of the cuticular layer of the spinning duct at different time points during silk spinning and the pupal stages. CL, cuticular layer. Scale bar, 10 μm. **(B)** Morphological analysis of the ASG. Scale bar, 200 μm. **(C)** Immunofluorescence analysis of ASSCP1, ASSCP2, and chitin in ASG. Blue, the nucleus; Red, ASSCP1 and ASSCP2 proteins; Green, chitin. Scale bar, 10 μm.

At the pre-pupation stage, some fissures could be observed on the cuticular layer. At the pupal stage, the cuticular layer was completely degraded and disappeared.

### Fine structure of the cuticular layer in silkworm spinning duct

TEM was used to observe the ultrastructure of the cuticular layer. It can be seen from Fig. 5 that the chitin in the cuticular layer constitutes two-dimensional horizontal sheets, the laminae. The cross-sectional diagram shows that the laminae had a network structure formed by the crosslink of chitin micro-filaments (Fig. 5A). Gaps (∼2 μm in diameter) between the chitin filaments were found, which may facilitate the transportation of water, ions, and other small molecules. From the longitudinal section shown in Fig. 5B, the laminae stacked together to form a multi-layer structure, with an interlayer spacing of ∼500 nm. The orientation of the cuticular layer is the same as the direction of silk protein transportation. Furthermore, immunoelectron microscopy (IEM) was introduced to study the localization of cuticle protein in the cuticular layer. The colloidal gold particles in Fig. 5C represent the cuticle protein ASSCP1. It can be seen that after being synthesized, ASSCP1 was stored in the secretory vesicles between the cuticular layer and gland cells, and then further secreted into the cuticular layer (Fig. 5C’). Magnifying the image found that ASSCP1 is mainly distributed in the interspace between laminae (Fig. 5C’’).

**Fig. 5.**
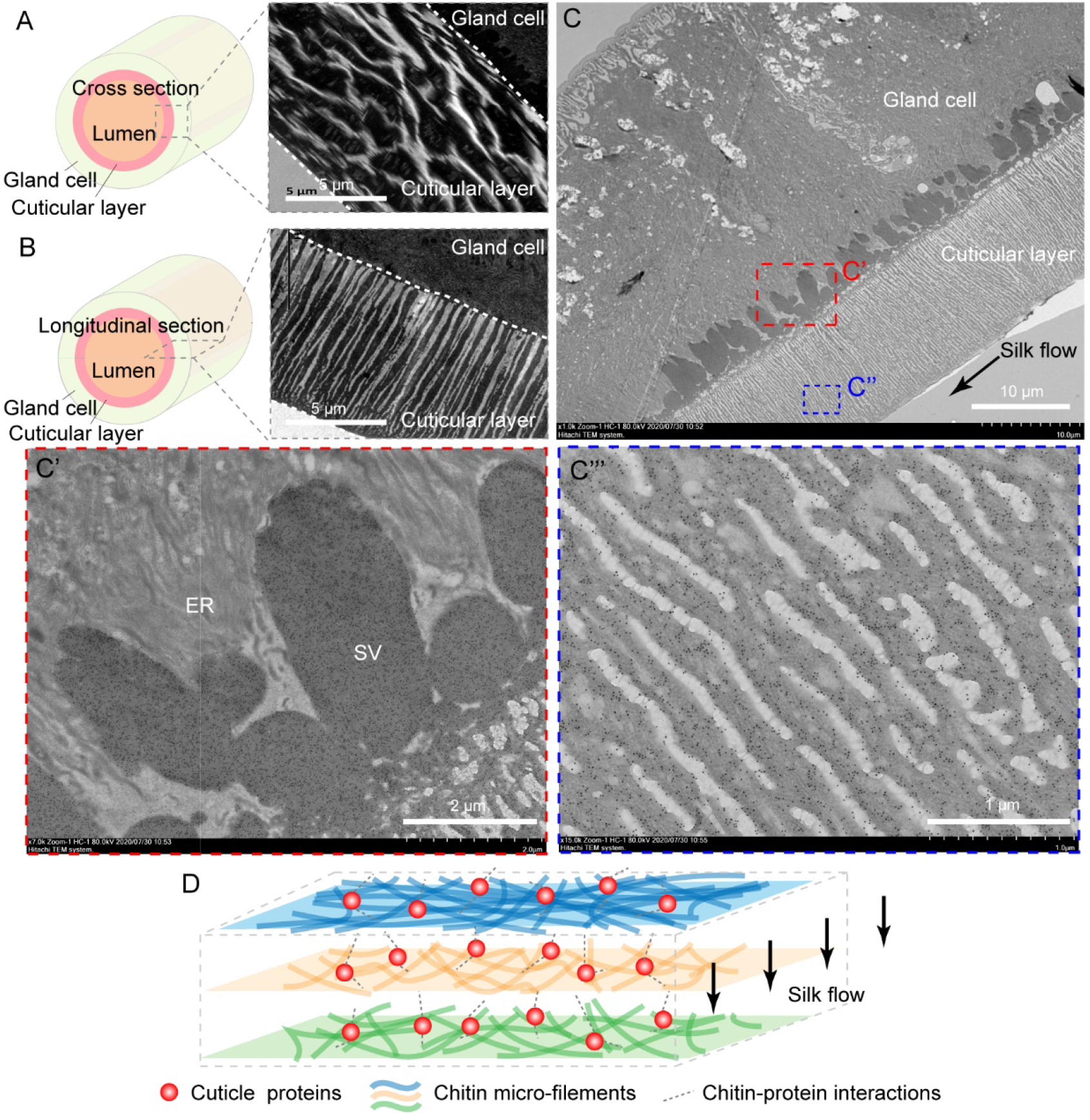
Fine structure of the cuticular layer of silkworm spinning duct. **(A)** TEM images of the cross-section of the cuticular layer. **(B)** TEM images of the longitudinal section of the cuticular layer. **(C)** Immunoelectron microscopy shows the localization of ASSCP1 protein, which is marked as black dots. Insets of C’ and C’’ show the distributions of ASSCP1 in the secretory vesicles (C’) and cuticular layer (C’’), respectively. The arrow indicates the flow direction of silk protein. ER, endoplasmic reticulum; SV, secretory vesicle. **(D)** Schematic illustration of the fine structure of the cuticular layer.

A schematic illustration of the ultrastructure of the cuticular layer is shown in Fig. 5D. The cuticular layer of the silkworm spinning duct is a multilayer structure by layered stacking of the chitin laminae. Inter- or intra-laminar interactions between cuticle proteins and chitin micro-filaments may stabilize the fine structure of the cuticular layer.

### Knocking down ASSCP1 and ASSCP2 expressions affect the size of ASG and disrupt the fine structure of the cuticular layer

To investigate the functions of cuticle proteins and chitin in the spinning duct, we successfully constructed two transgenic silkworm lines (*ASSCP1*^*-*^ *and ASSCP2*^*-*^) in which the expression of ASSCP1 or ASSCP2 was knocked down, respectively (Figs. S5). Through cross-breeding, we obtained a transgenic line (*ASSCP1*^*-*^*/ASSCP2*^*-*^) that both ASSCP1 and ASSCP2 were knocked down. The economic traits of the transgenic animals from all three lines remained unchanged, and no obvious changes could be observed in the morphologies of the cocoons and pupae (Fig. S5). Molecular analysis showed that the expression of ASSCP1 or ASSCP2 was successfully knocked down from each corresponding silkworm line (Fig. S6). It can be also seen that both ASSCP1 and ASSCP2 were knocked down from the *ASSCP1*^*-*^*/ASSCP2*^*-*^ line (Fig. S6). The chitin content was also measured, and no significant difference was observed between the transgenic silkworm line and wild-type (*WT*) silkworms (Fig. S7). The chitin content was not changed implied that chitin synthesis and metabolism pathways were not affected after knocking down.

The most exciting finding is that knocking down the expressions of cuticle proteins affected the length of the spinning duct and the structural integrity of the cuticular layer (Fig. 6). The average lengths of the spinning duct in the *ASSCP1*^*-*^ and *ASSCP1*^*-*^*/ASSCP2*^*-*^ transgenic lines decreased significantly from 23.60 mm (*WT*) to 20.75 mm and 21.05 mm, respectively (Fig. 6A and 6B). Observations showed that the decrease in length of the spinning duct in the transgenic line resulted from smaller gland cells (Fig. 6C). The small size of the gland cell did not cause any reduction in the diameter of the ASG (Fig. 6D). However, the thickness of the cuticular layer decreased dramatically (Fig. 6E). Knocking down the expression of ASSCP2 had a greater impact on the cuticular layer than ASSCP1, and a certain synergistic effect could be found when knocking down both two proteins. We further observed the ASG samples from different spinning periods and measured them at different positions (anterior, middle, and posterior) of the ASGs. The same results could be found (Fig. S8). To have a clear view of the changes in the cuticular layer, TEM analysis was performed. Compared with those from *WT* animals, the fine structure of the cuticular layer from the transgenic lines was disrupted. Cross-sections of the cuticular layer showed that chitin microfilaments became thinner, and gaps between chitin microfilaments within lamina were compressed (Fig. 6F). As listed in Table 1, the average width of the gaps decreased almost 3 times in the *ASSCP1*^*-*^*/ASSCP2*^*-*^ transgenic line, compared with those of *WT*. Longitudinal sections give more details of the changes (Fig. 6G). The average length of chitin laminae dropped to ∼2 μm (Table 1), suggesting that the chitin networks were interrupted by the gaps. The inter-space between the chitin laminae was larger in the transgenic lines, especially for *ASSCP1*^*-*^ and *ASSCP1*^*-*^*/ASSCP2*^*-*^ transgenic lines, compared with the *WT* (Table 1). Moreover, some unidentified substances filled the space between the chitin laminae. We suspect that one of the unidentified substances might be the unassembled chitin because there are insufficient cuticle proteins to bind to chitin.

**Fig. 6.**
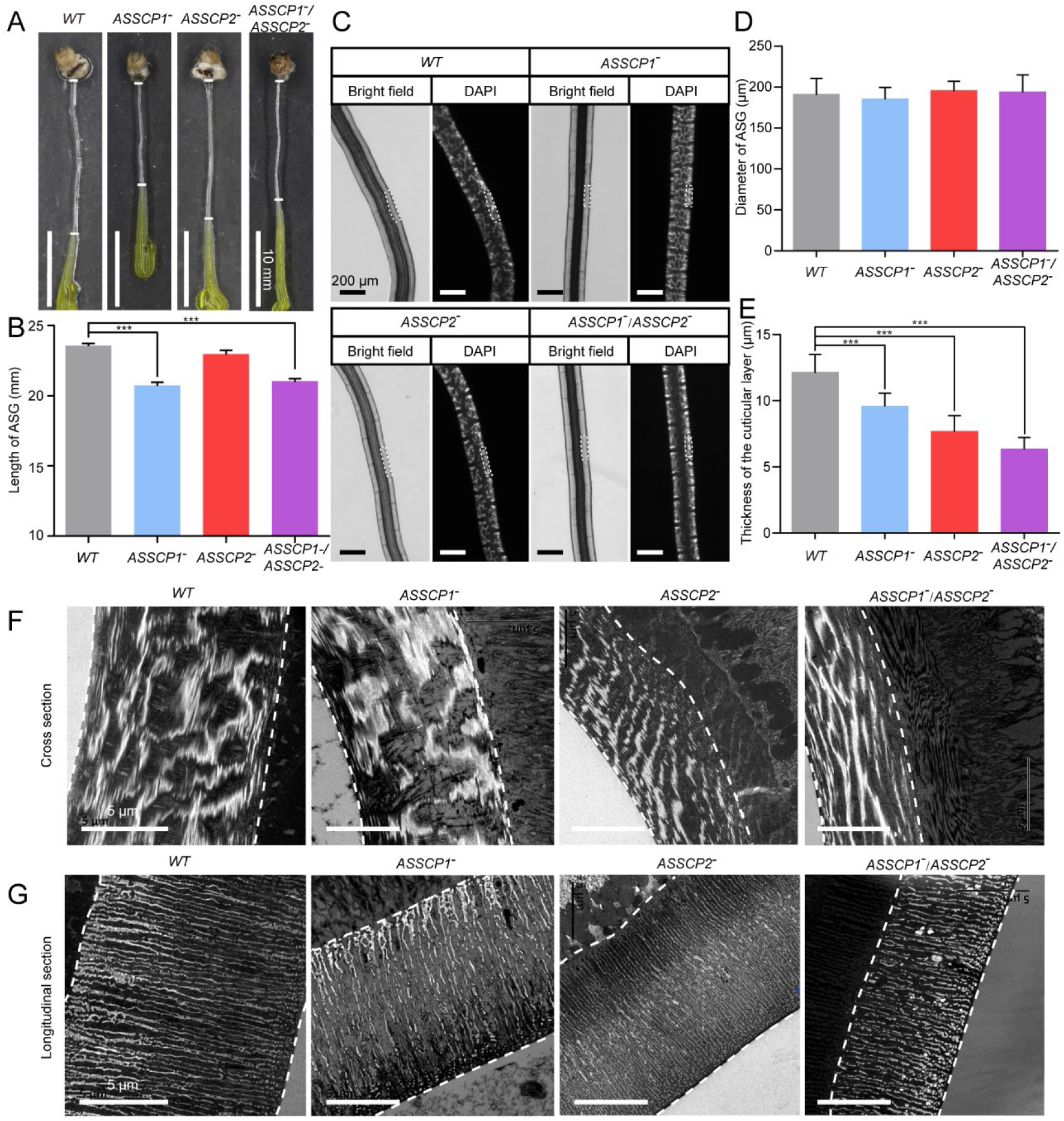
Knocking down the expressions of cuticle proteins affects the length and cell size of ASG and disrupts the fine structure of the cuticular layer. **(A)** Picture of the ASG from the transgenic and *WT* animals. The narrow duct between the two white lines is the ASG. Scale bar, 10 mm. **(B)** Comparison of the length of ASG from all silkworm lines. **(C)** Cell size decreased after knocking down ASSCP1 expression. Scale bar, 200 μm. The dotted box shows the single gland cell. **(D)** Comparison of the cross-sectional diameters of ASG from all silkworm lines. **(E)** Comparison of the thickness of the cuticular layer from all silkworm lines. Scale bar, 5 μm. **(F)** TEM images of the cross-section of the cuticular layer from all silkworm lines. **(G)** TEM images of the longitudinal section of the cuticular layer from all silkworm lines. Scale bar, 5 μm. ***, *p* < 0.001.

**Table 1.** Parameters for the structural changes of the cuticular layer after knocking down the expressions of ASSCP1 and ASSCP2.

| Parameter | WT | ASSCP1<sup>-</sup> | ASSCP2<sup>-</sup> | ASSCP1<sup>-</sup>/<br>ASSCP2<sup>-</sup> |
| --- | --- | --- | --- | --- |
| Width of the gaps between chitin microfilaments in the cross section ($\mu\text{m}$) | $1.77 \pm 0.94$ | $2.19 \pm 1.35$ | $0.95 \pm 0.33$ | $0.61 \pm 0.17$ |
| Length of the chitin laminae ($\mu\text{m}$) | $7.67 \pm 1.55$ | $2.69 \pm 1.21$ | $2.56 \pm 1.12$ | $1.73 \pm 0.56$ |
| Width between adjacent chitin laminae ($\mu\text{m}$) | $0.28 \pm 0.09$ | $0.59 \pm 0.09$ | $0.29 \pm 0.05$ | $0.53 \pm 0.16$ |

### Knocking down the expression of cuticle protein affects silk fibrillogenesis and fiber mechanical properties

We found that the fine structure of the cuticular layer was disrupted after knocking down the expressions of ASSCP1 and ASSCP2. The relationship between the spinning duct and silk fibrillogenesis prompted us to further investigate the characteristics of silk produced by these transgenic silkworms. SEM images show that the morphologies of the cocoon surface (Fig. S9) and the single silk fiber (Fig. 7A) were similar between the transgenic and *WT* silkworms. However, the diameter of the single silk fiber from the transgenic lines slightly decreased (Fig. 7C). Further, polarized light microscopy was used to compare the mesostructures of silk fibers (Fig. 7B). The brilliant color under cross-polarized light confirmed the high orientation of all kinds of fibers. However, a deep comparison of the images in Fig. 7B shows that there were several defects (white arrowheads) on the surface of the fiber from the knocking-down silkworms, whereas only a uniform and smooth morphology was observed on the surface of the *WT* silk fibers. A possible explanation of this observation would be that the disruption of the fine structure of the cuticular layer affected silk self-assembly and orientation during silk fiber formation.

**Fig. 7.**
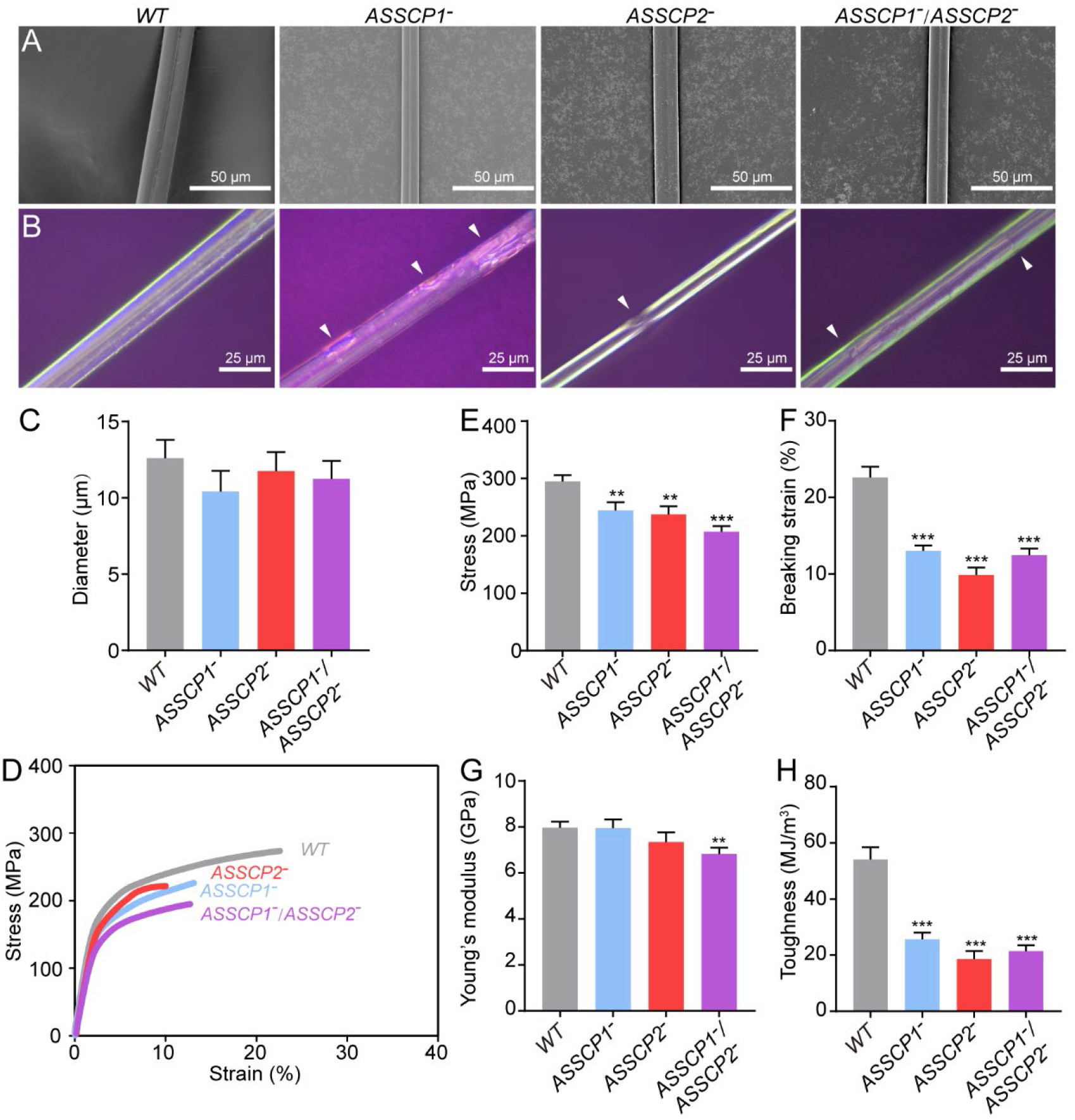
Knocking down the cuticle proteins affects silk fibrillogenesis and fiber mechanical properties. **(A)** SEM images of the single silk fiber. **(B)** Polarized light microscopy images of the single silk fiber. The arrowheads indicate the defects on silk fiber. **(C)** Diameters of the single silk fibers from different strains of the silkworm. **(D)** Averaged stress-strain curves of the single silk fibers. **(E)** Breaking stress of the single silk fibers. **(F)** Breaking strain of single silk fibers. **(G)** Young’s modulus of single silk fibers. **(H)** Toughness of single silk fibers. **, *p* < 0.01. ***, *p* < 0.001.

We introduced FTIR analysis to quantify the content of each secondary structure of silk fibers. Amide I band was used to perform the peak deconvolution. Then, we determined the content of each secondary structure by measuring the ratios of areas under the Gaussian peaks indicative of different secondary structures (Fig. S10). Table 2 lists the content of each secondary structure of silk fiber from all silkworm lines. We found that the content of β-sheets decreased in the knocking down silkworm strains, while the content of coils and helices, and β-turn structure increased. The results obtained by morphological and structural characterizations indicate that knocking down the expression of cuticle protein affects silk fibrillogenesis.

**Table 2.** Secondary structures of silk fibers.

| Silkworm strains | β-sheets | Coils & Helices | β-turn |
| --- | --- | --- | --- |
| *WT* | 44.99 ± 1.30 | 15.13 ± 0.83 | 9.46 ± 0.64 |
| *ASSCP1*<sup>-</sup> | 39.36 ± 0.93<sup>a</sup> | 21.44 ± 0.24<sup>b</sup> | 18.54 ± 0.56<sup>c</sup> |
| *ASSCP2*<sup>-</sup> | 39.28 ± 1.76 | 21.40 ± 0.55<sup>b</sup> | 18.58 ± 1.11<sup>c</sup> |
| *ASSCP1*<sup>-</sup>/*ASSCP2*<sup>-</sup> | 38.69 ± 0.13<sup>a</sup> | 21.21 ± 0.10<sup>b</sup> | 18.80 ± 0.10<sup>c</sup> |

a, b, and c represent the statistical differences in the corresponding secondary structures between the transgenic and wild type strains (Student's *t*-test).

To determine whether these structural changes have negative impacts on the mechanical properties of silk fiber, we carried out tensile tests (Figs. 7D and S10). The averaged stress-strain curves of both kinds of fibers are shown in Fig. 7D, and it can be seen that the fibers from the transgenic silkworms exhibited reduced breaking stress, breaking strain, and toughness (the area of the curve). Still, a certain synergistic effect could be found when knocking down both two proteins. Comparisons of mechanical parameters such as stress, strain, Young’s modulus, and toughness are also shown in Fig. 7. These data suggest that knocking down the expression of *ASSCP1* or/and *ASSCP2* had negative impacts on the mechanical properties of silk fiber, especially on silk extensibility and toughness.

## DISCUSSION

In this study, we identified chitin in the cuticular layer of the spinning duct of silkworms, a representative of silk-spinning arthropods. Using functional studies of the chitin-binding proteins ASSCP1 and ASSCP2, we show that chitin, ASSCP1, and ASSCP2 are the main components of the cuticular layer of the silkworm spinning duct. They participate in silk fibrillogenesis and regulate the final mechanical properties of the silk fiber.

Cuticle proteins are structural proteins, which can form the arthropod exoskeletons with chitin (Moussian, 2013). Cuticle proteins can be divided into several families, such as CPR (Karouzou et al., 2007), CPF/CPFL (Togawa, Augustine Dunn, Emmons, & Willis, 2007), CPT (Guan, Middlebrooks, Alexander, & Wasserman, 2006), CPG (Futahashi et al., 2008), CPAP1/CPAP3 (Behr & Hoch, 2005), and CPLC (Cornman & Willis, 2009). Among these families, the CPR family is the most widely distributed and found in *Lepidoptera, Hymenoptera, Coleoptera, Orthoptera, Diptera*, and *Hemiptera* (Willis & Iconomidou, 2005). The most obvious feature of the CPR family is the chitin-binding consensus, which is called the R&R consensus (Iconomidou, Willis, & Hamodrakas, 2005). The CPR family can be further divided into three types, RR-1, RR-2, and RR-3 (Andersen, 2000). Both ASSCP1 and ASSCP2 were identified as RR-2 cuticle proteins (Wang, Li, Liu, Xia, et al., 2017; Yi et al., 2013). The RR-2 consensus is rich in histidine, and the lysine residue is very conserved. These residues have been found to act as the reaction site for chitin-binding and cuticle sclerosis (Kerwin et al., 1999; Rebers & Willis, 2001). Thus, the RR-2 type cuticle proteins are generally found in hard cuticles (Willis, 2010). From our TEM pictures, it can be seen that the chitin in the cuticular layer shares the same multi-layer structure as the chitin found in hard cuticles (Fig. 5), such as the exoskeleton (Neville, 1975). Based on this evidence, we concluded that the cuticular layer, which is composed of chitin and RR-2 type cuticle proteins, is hard and rigid.

In arthropods, chitin is present in tissues such as exoskeletons and peritrophic matrix. In recent studies, chitin has also been found in spinning ducts of silk-spinning arthropods such as spiders, silkworms, and caddisworms (Ashton & Stewart, 2019; Davies et al., 2013). This finding suggests that silk-spinning arthropods may have evolutionary homology. However, we are mainly interested in why the spinning duct of arthropods needs chitin. This study attempts to clarify the biological significance of chitin in the spinning duct of arthropods.

Studies have shown that silk proteins undergo a transition from liquid to a liquid crystal state in the spinning duct (Jin & Kaplan, 2003; Kerkam, Viney, Kaplan, & Lombardi, 1991). When liquid silk enters the spinning duct, the shear rate of the silk protein increases rapidly as the diameter of the spinning duct becomes extremely small (reduced from approximately 420 μm to 80 μm) (Asakura et al., 2007). The average shear rate of liquid silk fibroin is 0.1 to 0.8 sec^-1^ in the posterior silk gland, approximately 10^−3^ sec^-1^ in the middle silk gland, and 20 to 400 sec^-1^ in the anterior silk gland (Kataoka & Uematsu, 1977).

Therefore, silk proteins bear tremendous shearing and extensional stresses when they flow through the spinning duct. Shearing and extensional stresses are key factors in regulating silk fibrillogenesis. A large number of studies have shown that silk proteins can aggregate, self-assemble, and go through fibrillogenesis under shearing and extensional stresses (Eisoldt et al., 2010; Hagn et al., 2010; Holland et al., 2012; Leclerc, Lefevre, Gauthier, Gagne, & Auger, 2013; Rammensee et al., 2008). These stresses in turn induce the stretch and orientation of the silk molecular chains along the fiber axis (Asakura et al., 2007; Vollrath & Knight, 1999). In this step, the conformational transition of silk proteins from random-coil and helix-like conformations to mainly β-sheet-rich structures is promoted (Vollrath & Knight, 2001). Furthermore, the silk protein is converted into a solid fiber under a suitable pH gradient and metal ion strength and pulled out of the spinneret by head swing (Andersson, Johansson, & Rising, 2016; Sparkes & Holland, 2017; Wang, Li, Liu, Chen, et al., 2017).

Silk proteins have a very high shear rate when flowing through the spinning duct. It is easy to see that both the silk proteins and the wall of the spinning duct are subject to considerable shearing stress. The flow of the viscous silk protein fluid in the lumen of the hard and rigid cuticular layer is the major source of this large shear force, which promotes silk fibrillogenesis. On the other hand, because the wall of the spinning duct is a single cell structure, it is the hard cuticle of the inner wall of the duct lumen that bears the tremendous shearing force generated by the flow to avoid the destruction of gland cells. We have also observed a slight decrease in the thickness of the cuticular layer of the spinning duct during the silk spinning process (Fig. S8), suggesting that the cuticular layer may be worn down. This observation further proves this hypothesis. Thus, we suggest that the biological function of chitin in the cuticular layer of the spinning duct is to provide shearing stress during silk fibrillogenesis and protect gland cells from shear damage.

Ultrastructural analysis revealed that the cuticular layer is a multilayer structure by the stacking of chitin laminae (Fig. 5). The fine structure of the cuticular layer is crucial for maintaining the physiological function of ASG. The chitin laminae stack vertically along the duct and perpendicular to the direction of silk protein transport, ensuring that the shearing force generated during the flow of silk protein will not cause relative slippage between the laminae, thus improving the structural stability and strength of the ASG. Further, gaps between the chitin filaments within the lamina and interlayer spaces were found in the cuticular layer. We have mentioned above that pH, ion strength and water content also contribute to silk fibrillogenesis. Protons, metal ions and water molecules are transported across the gland cells and silk proteins during silk fibrillogenesis. The gaps and interlayer spaces facilitate the transportation of these molecules and ensure the proper pH, ion strength, and water content for silk fibrillogenesis in the ASG lumen.

We also found that the cuticular layer of the spinning duct is periodically degraded and reformed during the molting period of silkworm larvae (Fig. 3). Why does the cuticular layer need to be periodically reconstructed?

First, this phenomenon can ensure the normal growth and development of the spinning duct. The periodic reconstruction of the cuticular layer is similar to the periodic reconstruction of the arthropod exoskeleton. The exoskeleton and cuticular layer are both relatively hard. When the growth of soft and flexible tissue exceeds the support of the rigid outer matrix, degradation of the old matrix and formation of a new matrix occurs. Our observations indicate that, after the period of molting, the diameter of the silkworm spinning duct increases (Fig. S12). The duct reaches its maximum size when spinning silk, which is about twice compared with that of the 4^th^ molting.

Second, the periodic reconstruction of the cuticular layer ensures that the larvae of each instar can spin the silk fiber with good mechanical properties. In addition to spinning silk during the metamorphosis period, silkworms also spin silk at other larval developmental stages. At present, 8 kinds of silk have been identified in the silkworm larval stage (Dong et al., 2013; Peng et al., 2019). These types of silk keep the immobilized molting larvae from being blown away by heavy wind (Peng et al., 2019) and facilitate molting by anchoring their ventral feet and the old cuticle to the substratum (Hu et al., 2016).

In conclusion, this study clarified the function of chitin in the spinning duct of silk-spinning arthropods. We found that the cuticle proteins ASSCP1 and ASSCP2 are co-located and co-expressed with chitin in the cuticular layer of the spinning duct. The cuticle proteins, together with chitin, assemble in the molting stage and degrades during spinning and metamorphosis. We conclude that these cuticle proteins can bind chitin and form the hard and rigid cuticular layer for supporting the cells and contributing to silk fibrillogenesis and spinning. Thus, these proteins may be the potential targets for improving the mechanical properties of silk fiber in the future. Our next works could overexpress these cuticle proteins in the spinning duct to produce high-performance silk fibers.

## AUTHOR CONTRIBUTIONS

**X. Wang:** Conceptualization, Methodology, Writing - Original Draft, Visualization, Funding acquisition. **X. Xie:** Methodology, Formal analysis, Investigation, Visualization. **K. Xie:** Formal analysis, Investigation. **Q. Liu:** Data curation. **Y. Li:** Data curation, Writing - Review & Editing. **X. Tan:** Validation. **H. Dong:** Resources. **X. Li:** Validation. **Z. Dong:** Writing - Review & Editing. **Q. Xia:** Conceptualization, Supervision. **P. Zhao:** Writing - Review & Editing. Project administration, Funding acquisition.

## ACKNOWLEDGEMENTS

This work was supported by grants from National Natural Science Foundation (Grants No. 31902216 and 31530071) and Natural Science Foundation of Chongqing, China (Grant No. cstc2020jcyj-cxttX0001).

