## Supplemental Figures 1-12 and Supplemental Table 1-2 for "Chitin and cuticle proteins form the cuticular layer in spinning duct of silk-spinning arthropods"

1 10 20 30
M I V N I F V A V F I L G V S A I N V V T G A S F S K V F V

31 40 50 60
Q G G R E V V P V E Y L Q Y G A P S I A V A G S Q L A Q I S

61 70 80 90
A A P V G S V V P N L V A P C G T P C I Y P G Q S I A P A T

91 100 110 120
T N V V T Y K S S V P V V V T N K E D A **A G Y E Y S Y L V Y**

121 130 140 150
**D E N T G D H K T Q H E L S D G F N A V V K K F L** P V N V E

151 160 170 180
K K I E Q H E K S H P D P P C H E V K N E Q L K V E T K T E

181 190 200 210
H S E S H H A P I V E E H H E T K S E E S S E S S S E E S H

211 220 230 240
E E K P T E N N H E E K E S S E S E S T E E S H E E K H S E

241 250 260 270
E A H H E V V A E A P H E E H S E P L K E L S T E S H E S S

271 280 290 300
S E E H S A E E H H V E P S K H E A L H D A P A H I E H V E

301 310 320 330
E T S H E E E K H E T S P H D A E E H T H E H T E T E T H E

331 340 350 360
H I S I N E A E H N A Q P Q Y N D I L K C V N A A I N T A A

361 370
G V A P Q R S E S P L T Y I I L N K P C

**Fig. S1** Sequence of silkworm ASSCP1 protein. The sequence in blue was selected to generate the recombinant proteins in *E. coli*. The red sequence represents the RR-2 domain which is predicted to have the chitin-binding activity.

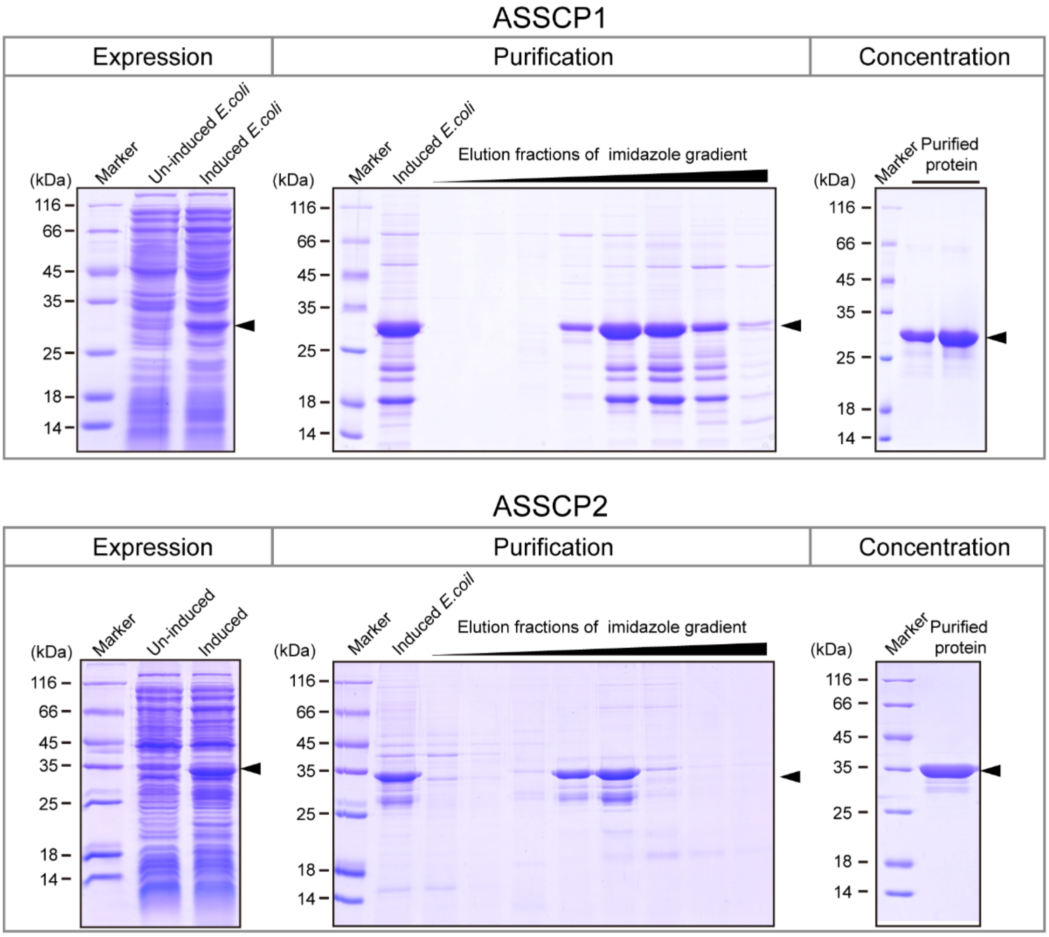


**Fig. S2** Expression and purification of recombinant proteins. Recombinant proteins were expressed and purified using Ni-NTA affinity chromatography and a stepwise imidazole gradient. Then, the recombinant proteins were concentrated. The arrows indicate the target proteins. Gel condition, 12 % SDS-PAGE followed by Coomassie brilliant blue G-250 staining.

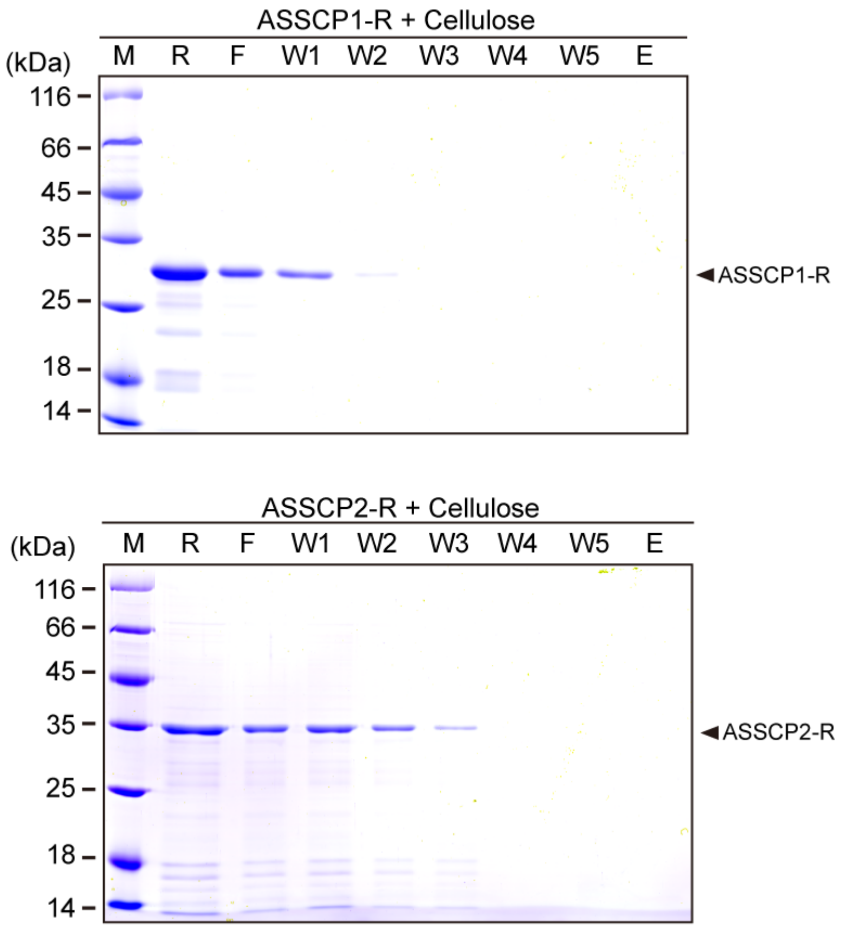


**Fig. S3** Cellulose binding assay of the recombinant proteins. No positive band can be detected in the eluted fraction, indicating the recombinant proteins are unable to bind with the cellulose. All fractions were detected using SDS-PAGE followed by Coomassie brilliant blue G-250 staining. M, protein marker; R, recombinant protein; F, flow-through fraction; W1-W5, washing fractions; E, eluted fraction.

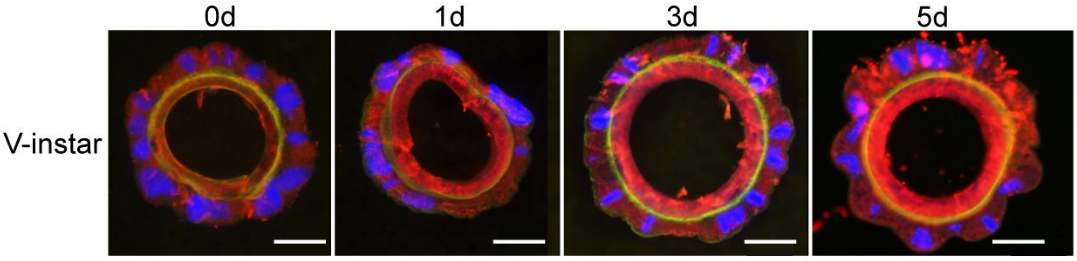


**Fig. S4** Immunofluorescence analysis of chitin and ASSCP1 in ASG during the fifth instar. Blue, nucleus; Green, chitin; Red, ASSCP1 protein; Scale bar, 50 μm.

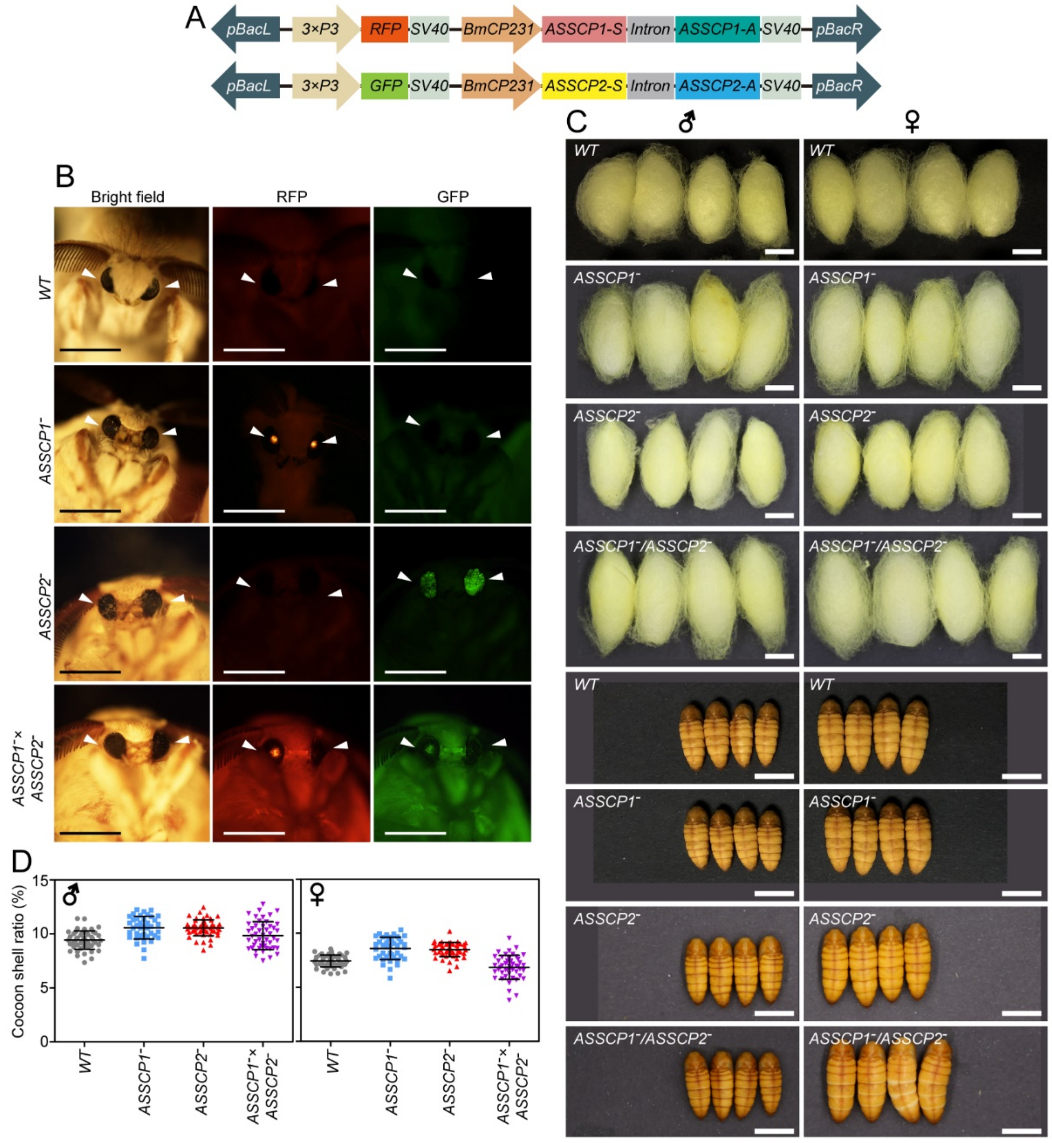


**Fig. S5** Transgenic silkworm acquisition. **(A)** Transgenic vector design. **(B)** Transgenic silkworms were isolated by detecting the RFP or GFP expression in the compound eyes. Scale bar, 10 mm. **(C)** Morphological observations of cocoons and pupae show no obvious changes between transgenic and wild-type silkworms. Scale bar, 10 mm. **(D)** Economic traits of silkworms were not changed after knocking down the expressions of ASSCP1 and ASSCP2.

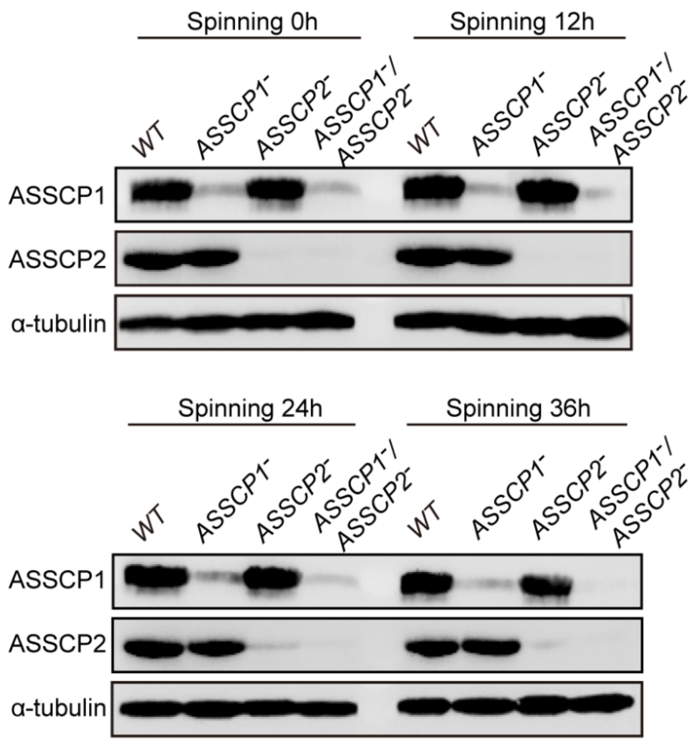


**Fig. S6** Molecular analysis of transgenic silkworm. Western blotting analysis shows that the expression of ASSCP1 protein or ASSCP2 protein was successfully knocked down in the corresponding transgenic line, respectively, during the spinning stage. Furthermore, both ASSCP1 and ASSCP2 were knocked down from the ASSCP1$^-$/ASSCP2$^-$ hybrid line. Silkworm α-tubulin protein was used as an internal control.

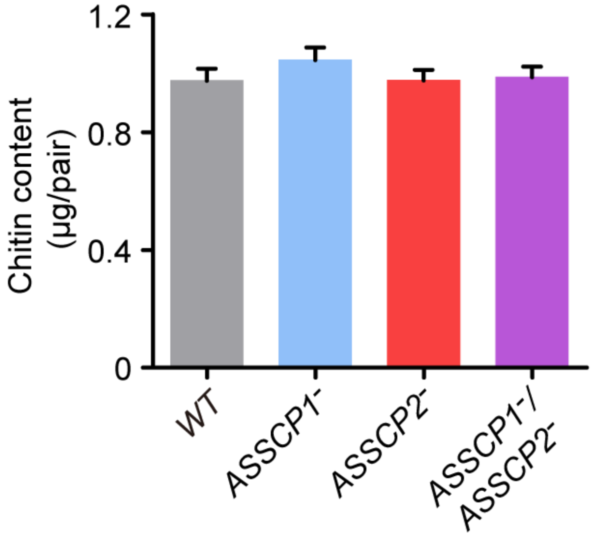


**Fig. S7** Chitin content of the ASG.

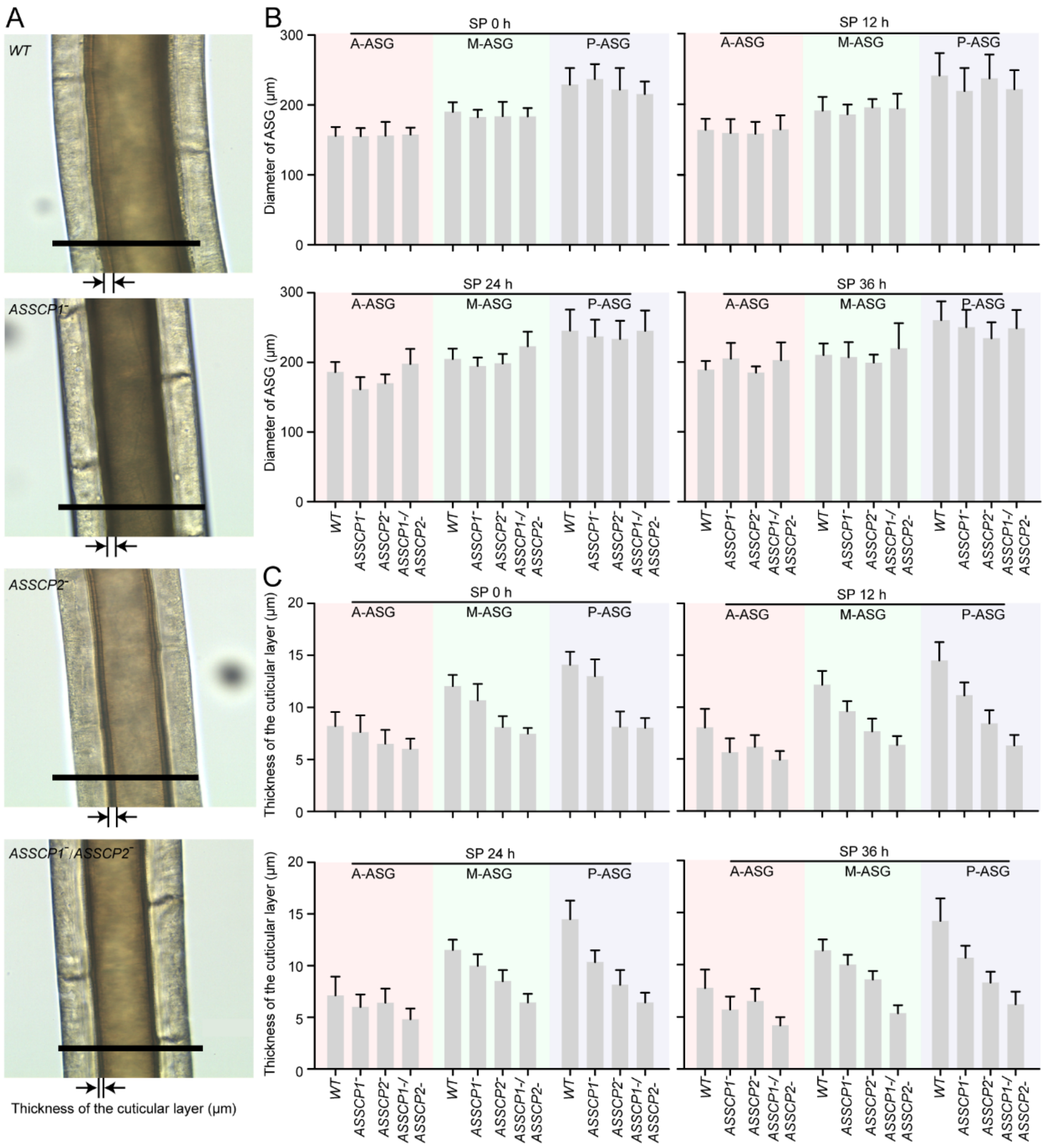


**Fig. S8** Morphological characteristics of the ASG of the transgenic silkworm lines. **(A)** Morphological observation of ASG using an optical microscope. Scale bar, 200 μm. (B) Comparisons of ASG diameter between all silkworm lines during the spinning stage. (C) Comparisons of the thickness of the cuticular layer of ASG between all silkworm lines during the spinning stage.

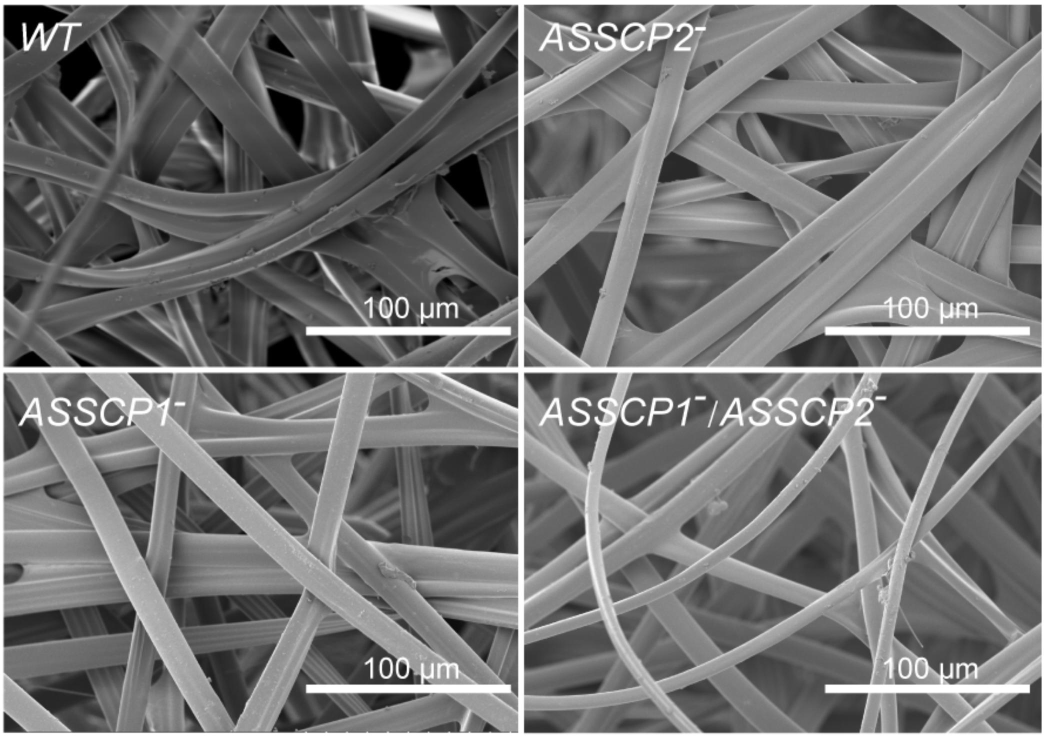


**Fig. S9** SEM analysis of the cocoons.

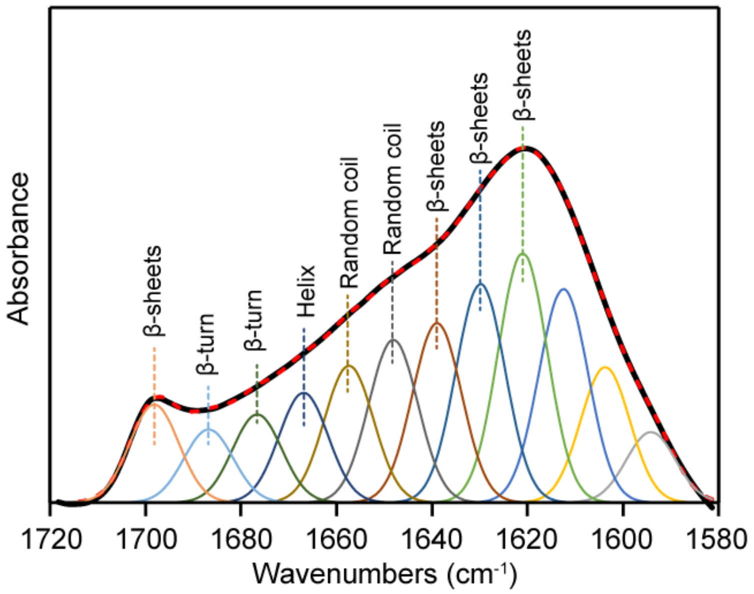


**Fig. S10** Deconvolution of amide I band of silk fibers. Black curve, original spectrum. Colorful curve, deconvoluted peaks indicative of different secondary structures. Red dashed curve, the simulated spectrum from summed peaks.

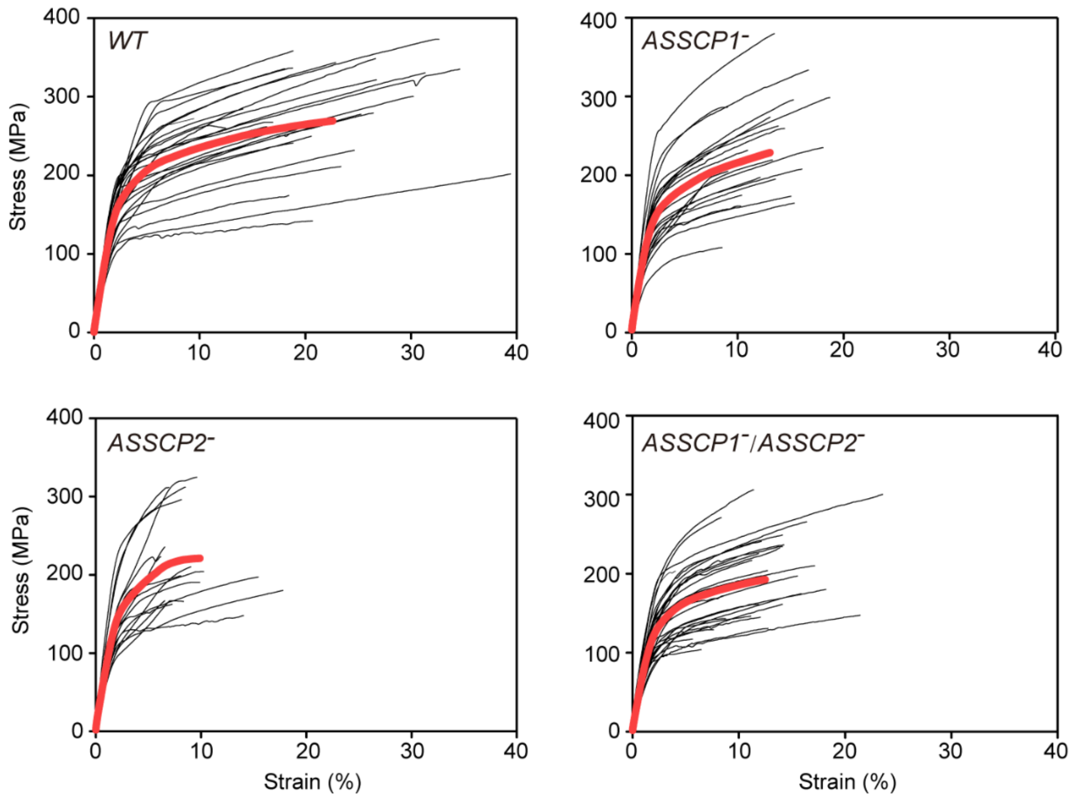


**Fig. S11** Stress-strain curves of the silk fibers. The red curve represents the averaged stress-strain curve of each group.

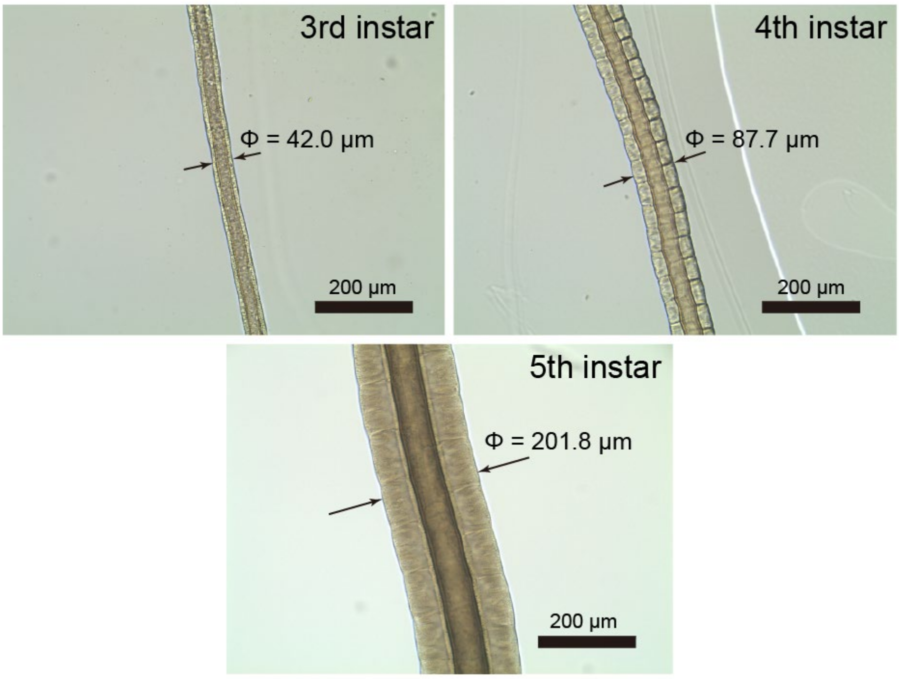


**Fig. S12** Spinning duct diameter increases 2-fold as the larvae enter a new instar. The diameter of the spinning duct from the 3$^{rd}$, 4$^{th}$, and 5$^{th}$ instar larvae is 42.0, 87.7, and 201.8 μm, respectively.

**Table S1.** Primers used in this study.

| Purpose | Primer name | Sequence (5' to 3') |
|---|---|---|
| Protein expression | *ASSCP1-F (Nde I)* | GGAATTCCATATGCAAGGCGGACGAGAAGTA |
| | *ASSCP1-R (Xho I)* | CCGCTCGAGTTAAGAACTTTCTTCACTTTTC |
| | *ASSCP2-F (EcoR I)* | CGCGAATTCGCAGCAGCTCCTGCAGTAGCGG |
| | *ASSCP2-R (Xho I)* | GCGCTCGAGTTAGGGGTAACGGTATCCAGCA |
| Transgenic vector construction | *BmCP231-F (Sal I)* | GCGTCGACGCTTCACGGCAGAAATAG |
| | *BmCP231-R (EcoR I)* | CGGAATTCTGCTGTGGTTGTTGAGCT |
| qRT-PCR | *ASSCP1-F* | TGTAACTGGTGCTTCATTCTCT |
| | *ASSCP1-R* | CGCGACTAGATTTGGTACTACT |
| | *ASSCP2-F* | TCAATGCCGTAGTACGTAAAGA |
| | *ASSCP2-R* | TATCTGGCTGCAGTAACATAGG |
| | *BmTif4a-F* | TTCGTACTGGCTCTTCTCGT |
| | *BmTif4a-R* | CAAAGTTGATAGCAATTCCCT |

Single underlined letters indicate restriction endonuclease cleavage sites.

**Table S2.** Sequence of dsRNAs used for transgenic vector construction.

| Gene fragement | Sequence (5'-3') |
|---|---|
| *ASSCP1 dsRNA* | GAATTCTTATCTTAGGCGTCTCCGCAATTAATGTTGTAACTGGTGCTTCA<br>TTCTCTAAAGTATTCGTACAAGGCGGACGAGAAGTAGTGCCAGTTGAAT<br>ACTTACAATATGGCGCTCCATCAATCGCCGTAGCAGGATCACAACTAGC<br>CCAAATATCTGCAGCCCCTGTAGGATCAGTAGTACCAAATCTAGTCGCG<br>CCATGTGGAACGCCCTGTATGTGAGCTCATCGATTCTGGACTATGCACT<br>TCGCCTCTCGGCCGGTGGGCCGTTATCGACCGTTATCTGACGAATGAC<br>TTTGTTCTGTTTCAGATACAGGGCGTTCCACATGGCGCGACTAGATTTG<br>GTACTACTGATCCTACAGGGGCTGCAGATATTTGGGCTAGTTGTGATCC<br>TGCTACGGCGATTGATGGAGCGCCATATTGTAAGTATTCAACTGGCACT<br>ACTTCTCGTCCGCCTTGTACGAATACTTTAGAGAATGAAGCACCAGTTA<br>CAACATTAATTGCGGAGACGCCTAAGATAATCTAGA |
| *ASSCP2 dsRNA* | ACGCGTAACGAGCGGGTACTACTGCGGGCTCTGCAACAACTGCAGGC<br>GCAGCCGCCACGAGTGGCTCTTTACGTACTACGGCATTGAAGCCATTA<br>ATGGAGTCAGCGGCGTAGTCCACGACACGACGTGTGCCGTCGGGTTC<br>AACCAAAGAGTAAGATCCTTGAACCAAATCTCCAGTGAGCTCATCGATT<br>CTGGACTATGCACTTCGCCTCTCGGCCGGTGGGCCGTTATCGACCGTT<br>ATCTGACGAATGACTTTGTTCTGTTTCAGTGGAGATTTGGTTCAAGGAT<br>CTTACTCTTTGGTTGAACCCGACGGCACACGTCGTGTCGTGGACTACG<br>CCGCTGACTCCATTAATGGCTTCAATGCCGTAGTACGTAAAGAGCCACT<br>CGTGGCGGCTGCGCCTGCAGTTGTTGCAGAGCCCGCAGTAGTACCCG<br>CTCGTTGCGGCCGC |

Single underlined letters indicate restriction endonuclease cleavage sites; double underlined letters indicate the intron sequences.
